# A safer fluorescent *in situ* hybridization protocol for cryosections

**DOI:** 10.1101/2025.05.25.655994

**Authors:** Akane Chihara, Rima Mizuno, Norie Kagawa, Ayuko Takayama, Akinori Okumura, Miyuki Suzuki, Yuki Shibata, Makoto Mochii, Hideyo Ohuchi, Keita Sato, Ken-ichi T Suzuki

**Affiliations:** Trans-Scale Biology Center, National Institute for Basic Biology, Okazaki, Aichi, 444-8585, Japan; Department of Basic Biology, The Graduate University for Advanced Studies, SOKENDAI, Okazaki, Aichi, 444-8585, Japan; Department of Life Science, Graduate School of Science, University of Hyogo, Ako-gun, Hyogo, 678-1297, Japan; Division of Biology and Biological Engineering, California Institute of Technology, Pasadena, California, 91125, United States; Department of Biology, Nippon Medical School, Musashino, Tokyo, 180-0023, Japan; Department of Cytology and Histology, Faculty of Medicine, Dentistry, and Pharmaceutical Sciences, Okayama University, Okayama, 700-8558, Japan

**Keywords:** in situ HCR, SABER-FISH, low-toxicity, cryosection, amphibians, medaka, mouse

## Abstract

Fluorescent *in situ* hybridization (FISH) enables highly sensitive, high-resolution detection of gene transcripts. Moreover, by employing multiple probes, this technique allows for multiplexed, simultaneous detection of distinct gene expression patterns spatiotemporally, making it a valuable spatial transcriptomics approach. Owing to these advantages, FISH techniques are rapidly being adopted across diverse areas of basic biology. However, conventional protocols often rely on volatile, toxic reagents such as formalin or methanol, posing potential health risks to researchers. Here, we present a safer protocol that replaces these chemicals with low-toxicity alternatives, without compromising the high detection sensitivity of FISH. We validated this protocol using both *in situ* hybridization chain reaction (HCR) and signal amplification by exchange reaction (SABER)-FISH in frozen sections of various model organisms, including mouse (*Mus musculus*), amphibians (*Xenopus laevis* and *Pleurodeles waltl*), and medaka (*Oryzias latipes*). Our results demonstrate successful multiplexed detection of morphogenetic and cell-type marker genes in these model animals using this safer protocol. The protocol has the additional advantage of requiring no proteolytic enzyme treatment, thus preserving tissue integrity. Furthermore, we show that this protocol is fully compatible with EGFP immunostaining, allowing for the simultaneous detection of mRNAs and reporter proteins in transgenic animals. This protocol retains the benefits of highly sensitive, multiplexed, and multimodal detection afforded by integrating *in situ* HCR and SABER-FISH with immunohistochemistry, while providing a safer option for researchers, thereby offering a valuable tool for basic biology.

## 1. Introduction

mRNA *in situ* hybridization (ISH) has become a critical method for studying the distribution of mRNA in tissues, organs, and even whole bodies in various organisms. Initially, ISH employed probes labeled with radioisotopes such as ^3^H and ^35^S, with detection achieved through autoradiography (Cox et al., 1984; Gall & Pardue, 1969; Singer & Ward, 1982). This was gradually superseded by a chromogenic detection method using probes labeled with biotin or digoxigenin to visualize target transcripts through enzymatic reactions using peroxidase or alkaline phosphatase (Tautz & Pfeifle, 1989). More recently, fluorescence *in situ* hybridization (FISH) has emerged, enabling imaging through fluorophore-labeled probes using fluorescence microscopy.

A transcript detection method, single-molecule FISH (smFISH), involves designing multiple specific-oligonucleotide probes complementary to particular target mRNAs, which hybridize to allow the detection of individual transcript molecules through fluorescence imaging (Raj et al., 2008). However, a simple increase in the number of probes alone is insufficient to achieve high sensitivity. To overcome this limitation, signal amplification techniques such as *in situ* hybridization chain reaction (*in situ* HCR) (Choi et al., 2010, 2018), RNAscope (Wang et al., 2012), and signal amplification by exchange reaction (SABER)-FISH (Kishi et al., 2019) have been widely adopted in the improved smFISH. *In situ* HCR amplifies the secondary probe signal through isothermal amplification without enzymatic reactions, while methods such as RNAscope and SABER-FISH utilize branched secondary probes to hybridize multiple detection (imager) probes with fluorophores, significantly enhancing signal strength. These techniques enable the simultaneous detection of several mRNAs using distinct fluorophore-labeled probes and allow multiplexed FISH by repeated rounds of stripping and re-hybridization of probes to detect multiple sets of targets on the same section. Consequently, these approaches facilitate the acquisition of the spatial transcriptomics data underlying numerous gene expression patterns from the same tissue sections or fixed samples, making them highly versatile and powerful techniques.

Like other histological methods, FISH requires tissue or organ fixation, lipid removal, and enhanced probe permeability. Aldehyde-based fixatives such as formalin and organic solvents such as methanol are widely used for these purposes in basic biological experiments. However, these reagents pose significant risks due to their toxicity and volatility, requiring researchers to handle them with great care to ensure worker safety. Given these safety concerns, it has become increasingly important to develop safer laboratory environments and protocols to protect the health of researchers. To address these issues, this study aimed to develop a FISH protocol using low-toxicity fixatives and organic solvents as a safer alternative to hazardous reagents such as formalin and methanol. The protocol was evaluated and optimized through experimental applications in basic biology focusing specifically on model animals such as *Mus musculus, Xenopus laevis*, *Pleurodeles waltl* and *Oryzias latipes*.

## 2. Result

### 2.1 Establishment of a safer FISH protocol

We developed a safer FISH protocol that does not rely on highly toxic chemicals and evaluated its suitability in various contexts in the investigations described below. The main features of this refined FISH protocol are as follows. First, when applied to mouse, amphibian, or medaka tissues, the protocol allows fresh tissues or organs to be immediately embedded and frozen in O.C.T compound without a pre-fixation step. This capability makes the protocol compatible with spatial RNA-seq (Chen et al., 2022; Ståhl et al., 2016) which often omits the pre-fixation step to ensure the high-quality mRNA sequences required for these spatial transcriptomics techniques. Similarly, pre-fixation is not necessarily required for FISH on cryosections. In our protocol, cryosections are post-fixed directly on the glass slides after sectioning. Generally, 2–3% glutaraldehyde, 4% paraformaldehyde (4% PFA), or 3.7% formaldehyde (10% formalin) is used for tissue fixation; however, in this study, we examined two commercially available fixatives: ALTFiX and PAXgene®. The former is glyoxal-based, a dialdehyde compound, while the latter is ethanol-based and contains proprietary components. Both reagents are stated by their manufacturers to have low toxicity.

Secondly, our FISH protocol employs ethanol instead of methanol for the fixation, delipidation, dehydration, and permeabilization processes in order to minimize toxicity. Tissue integrity was well preserved with either ALTFiX or PAXgene®, and robust fluorescence signals were successfully detected in *in situ* HCR and SABER-FISH (as described below). Although proteolytic enzymes such as proteinase K, pepsin, and trypsin have been widely used to enhance probe accessibility to target RNA in FISH to date, they can damage tissues or adversely affect the signal intensity of target mRNA if not carefully optimized. To circumvent this issue, our protocol does not include these proteolytic enzymes and instead uses a modified detergent solution containing Tween 20 and sodium dodecyl sulfate (SDS) for a 30-min incubation at room temperature (Bruce & Patel, 2022; Morabito et al., 2023). This treatment effectively provided strong fluorescence signals without causing substantial tissue damage or target RNA degradation. Importantly, this workflow also permits subsequent immunohistochemistry (IHC) staining to be performed directly after mRNA FISH without any additional blocking and antigen retrieval steps. As demonstrated below, antigenicity was sufficiently preserved to allow reliable antibody-based detection following *in situ* HCR, enabling combined visualization of mRNA transcripts and protein markers within the same tissue section. The workflow for this safer protocol is outlined in Figure 1.

**Figure 1.**
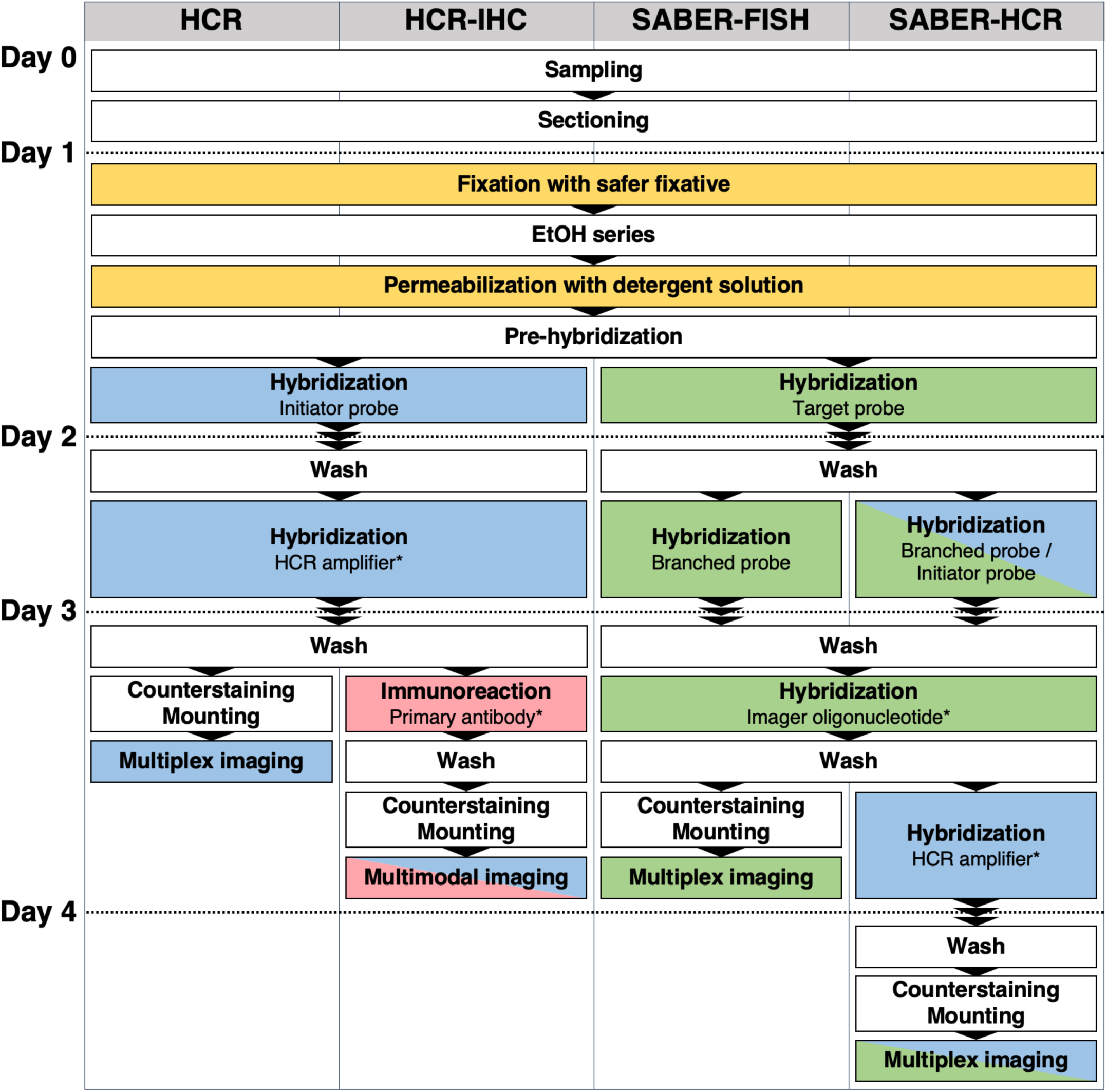
Schematic workflow of a safer FISH protocol. This figure illustrates the workflows for HCR, HCR-IHC, SABER-FISH, and SABER-HCR as utilized in this study. Tissues or organs are collected on Day 0, fixed if necessary, embedded in O.C.T compound, and then cryosectioned. On Day 1, the FISH protocol begins. Steps highlighted in yellow indicate the novel modifications introduced in this study. Cryosections mounted on glass slides are fixed with a safer fixative, dehydrated through a graded ethanol series, and then permeabilized with a detergent solution to enhance probe penetration. Subsequent steps follow the standard HCR or SABER-FISH workflow thereafter. Steps highlighted in blue and green indicate the HCR steps and SABER-FISH steps, respectively. In this study, the SABER-FISH branching step (indicated as “Hybridization branched probe” in this figure) was performed once; however, it can be repeated multiple times if further amplification is required. Steps highlighted in red indicate immunohistochemistry (IHC) steps on Day 3. In the HCR-IHC experiments conducted here, protein detection via IHC was performed following mRNA detection by FISH. Asterisks indicate that the HCR amplifier, primary antibody, and imager oligonucleotide are fluorescently labeled.

### 2.2 Application of the safer protocol for multiplexed detection of limb development-related genes in mouse limb buds

To evaluate the applicability of our safer protocol for multiplexed molecular detection in mammalian tissues, we performed *in situ* HCR on mouse limb bud sections. Cryosections were fixed using either conventional 4% PFA (Figure 2A–C) or the formaldehyde-free fixative ALTFiX (Figure 2D–F), followed by identical downstream processing. We simultaneously examined the expression of *Sox9* and *Col2a1*, both of which are key regulators of limb skeletal development. During limb development, *Sox9* encodes a master transcription factor for chondrogenic lineage specification and is required for the initiation of cartilage formation (Figure 2A, D), whereas *Col2a1* encodes a major cartilage extracellular matrix component expressed in differentiating chondrocytes (Figure 2B, E) (Akiyama et al., 2002; Bi et al., 1999). Consistent with their known biological roles, *in situ* HCR demonstrated spatially distinct yet partially overlapping expression domains of these genes within the developing mouse limb bud (Figure 2C, F). *Sox9* and *Col2a1* signals were prominently detected in regions corresponding to the cartilage primordia. Importantly, comparable signal patterns and fluorescence intensities were obtained under both PFA and ALTFiX fixation conditions, indicating that ALTFiX fixation supports multiplexed *in situ* HCR detection with performance equivalent to conventional PFA fixation. The preservation of nuclear architecture and overall tissue integrity under ALTFiX fixation is clearly evidenced by the DAPI images (Figure 2C, F).

**Figure 2.**
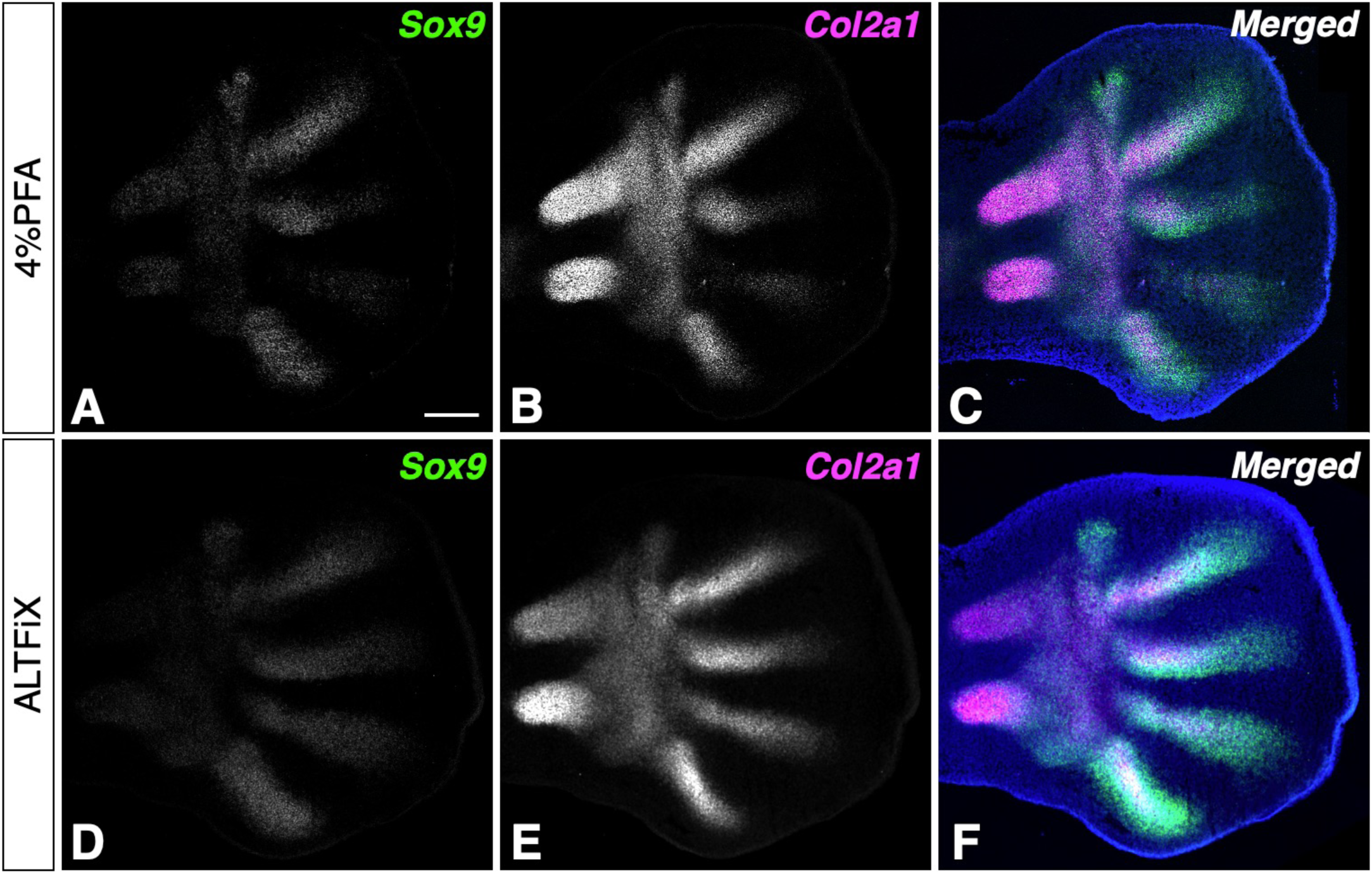
Comparison of multiplexed HCR-based mRNA detection in mouse limb buds using conventional 4% paraformaldehyde (PFA) and ALTFiX fixation. Representative images of mouse limb bud cryosections fixed with either (A–C) conventional PFA or (D–F) the formaldehyde-free fixative ALTFiX. Samples were processed in parallel under identical downstream conditions. (A, D) *Sox9* mRNA; (B, E) *Col2a1* mRNA; and (C, F) merged images with DAPI. Both *Sox9* and *Col2a1* signals were prominently localized to the cartilage primordia. The comparable signal patterns and fluorescence intensities between the two groups indicate that ALTFiX fixation supports multiplexed *in situ* HCR detection at levels equivalent to conventional PFA fixation. Nuclei were counterstained with DAPI. Scale bar: 200 µm.

### 2.3 Application of the safer protocol for multimodal detection in frogs

We next tested this protocol in a model amphibian, *X. laevis*, examining limb buds and adult elbow joints. In previous reports, *shh* is localized to the posterior mesenchyme, whereas *fgf8* is expressed in the apical ectodermal ridge (AER) in frog limb bud (Christen & Slack, 1997; Endo et al., 1997). As anticipated, *shh* and *fgf8* mRNAs were simultaneously detected in stage 52 tadpole limb buds (Figure 3A, B). In developing frog hindlimbs, *hoxa13* is expressed in the autopod region, while *msx1* is localized more distally, directly beneath the AER. Consistent with earlier studies (Barker & Beck, 2009; Endo et al., 2000), *in situ* HCR confirmed that *msx1* was expressed in the distal region of stage 53 hindlimb bud, whereas *hoxa13* signals were detected in the broader presumptive autopod region (Figure 3C, D). Finally, *in situ* HCR with our protocol specifically visualized the cartilage marker *aggrecan* (*acan*) and the elbow joint marker *proteoglycan4* (*prg4*) in adult forelimb elbow joints (Figure 3E) (Rhee et al., 2005; Watanabe et al., 1994).

**Figure 3.**
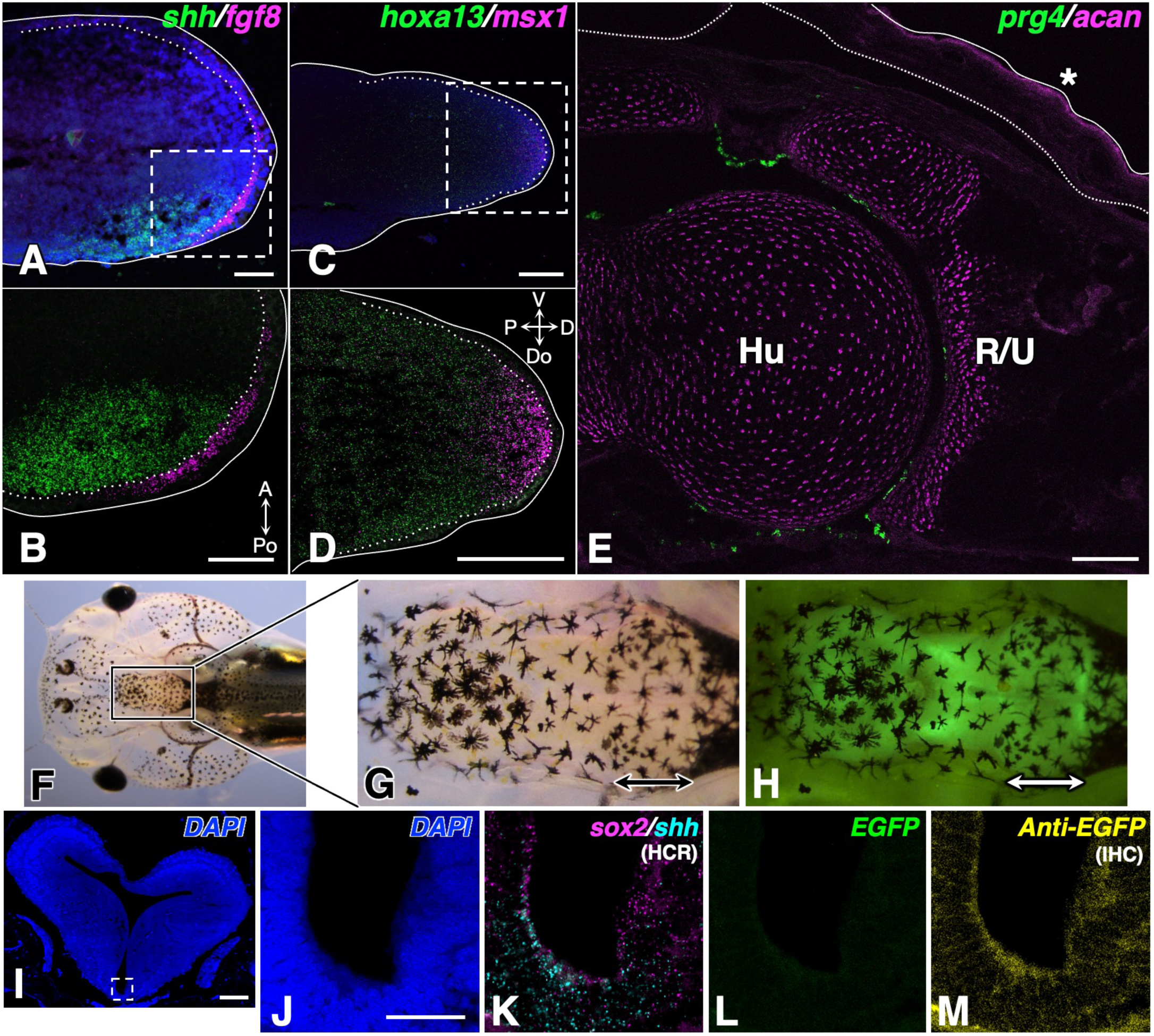
Multiplexed and multimodal detection by *in situ* HCR and its integration with IHC in frog tissue sections. (A–E) *In situ* HCR performed using the safer protocol in limb tissue sections from stage 52 to 53 tadpoles and adult frogs. (A, B) *shh* (green) and *fgf8* (magenta) mRNAs are localized to the posterior hindlimb bud mesenchyme and the AER, respectively. (A) Merged view of the three channels. (B) Magnified view of the region outlined by the dashed line in A. (C, D) Complementary expression of *hoxa13* (green) and *msx1* (magenta) exhibit along the proximal-distal axis in stage 53 hindlimb bud. (D) Magnified view of the region outlined with a dashed line in C. Nuclei were counterstained with DAPI in A and C. (E) *prg4* (green) and *acan* (magenta) signals in the articular cartilage and chondrocytes of the adult forelimb elbow joint, respectively. The asterisk marks autofluorescence in the outer layer of the epidermis. Sections in A and B were post-fixed with PAXgene®, whereas sections in C, D, and E were fixed with ALTFiX. Dotted white lines indicate the epithelial-mesenchymal boundary and solid white lines outline the limb buds or adult forelimb in A–E. (F–M) Combined *in situ* HCR and immunohistochemistry (HCR-IHC) in brain sections from stage 50 knock-in tadpole (*sox2:egfp*). (F) Representative image of a knock-in tadpole (*sox2:egfp*). The boxed area indicates the region shown in the close-up in G and H. (G) Bright-field image of the boxed region shown in F. (H) Fluorescence image of the same field of view shown in G. Arrows in G and H indicate the midbrain, from which the brain sections used for the combined HCR and IHC were obtained. (I) Low-magnification DAPI images of a brain section. (J–M) Enlarged views of the ventral region outlined by a dashed line in I, corresponding to the individual channels shown in J–M. (J) DAPI nuclear staining. (K) *In situ* HCR signals for *sox2* (magenta) and *shh* (cyan) mRNAs. (L) EGFP fluorescence derived from the *sox2:egfp* reporter. (M) IHC using Alexa 647 labeled anti-EGFP antibody. Abbreviations: R/U, radius/ulna; Hu, humerus; A, Anterior; Po, Posterior; Do, Dorsal; V, Ventral; P, Proximal; D, Distal direction. Scale bars: (A, B, E, I) 100 µm; (C, D) 200 µm; (J) 50 µm.

To investigate the compatibility of *in situ* HCR with IHC in amphibian tissues (HCR-IHC), we applied the optimized protocol to brain sections from stage 50 knock-in tadpole (*sox2:egfp*). These tadpoles express EGFP driven by the endogenous *sox2* promoter/enhancer (Figure 3F–M) (Mochii et al., 2024). *In situ* HCR was performed to detect *sox2* and *shh* transcripts, followed by IHC for the EGFP protein on the same sections. *sox2* is known to be expressed in neural tissues and plays an essential role in the neural differentiation of early frog neuroectoderm, consistent with its expression in proliferative neural progenitor domains including the ventricular zone (Kishi et al., 2000). Similarly, *shh* expression in frog embryos has been documented in ventral midline regions such as the prospective floor plate during early neurulation, where it contributes to dorsoventral patterning of the central nervous system (Domínguez et al., 2010). In our experiments, *in situ* HCR signals for *sox2* and *shh* mRNAs were detected in brain sections from the midbrain ventral region (Figure 3K). While EGFP fluorescence from the *sox2:egfp* reporter was barely detectable in the same sections (Figure 3L), whereas robust anti-EGFP antibody signals were obtained following IHC (Figure 3M), demonstrating that IHC can be effectively performed after *in situ* HCR even without any blocking and antigen retrieval steps. To confirm the specificity of this EGFP detection, we performed the same experimental procedure on stage 50 wild-type (WT) tadpoles as a negative control experiment. Expectedly, while multiplexed *in situ* HCR signals were consistently detected in WT sections, no anti-EGFP antibody signal was observed at all (Figure S1A–G).

As expected from previous work, *sox2* HCR signals were enriched in neural progenitor domains including the ventricular zone, and *shh* signals were concentrated in ventral midline areas consistent with floor plate expression (Figure 3K). However, *sox2* HCR signals were also observed outside of the classical ventricular zone, suggesting broader or more heterogeneous *sox2* transcript distribution at this developmental stage. Importantly, the localization of reporter-derived EGFP protein did not completely overlap with that of endogenous *sox2* mRNA (Figure 3K–M), indicative of subtle differences between mRNA and protein distributions.

Together, these results demonstrate that *in situ* HCR and IHC can be performed sequentially on tissue sections, enabling the simultaneous visualization of endogenous transcripts and reporter proteins within a single specimen with sufficient sensitivity for both detection modalities (Figure 3F–M).

### 2.4 Application of the safer protocol for multiplexed detection of limb development-related genes in newts

To evaluate the suitability of this safer protocol for detecting mRNA, we performed *in situ* HCR to visualize the expressions of genes related to limb development in the hindlimb bud of a model amphibian, *P. waltl*. Using our protocol, we examined the expression patterns of *Fgf8* and *Fgf10* (Figure 4A–D). These two factors are known to form a positive feedback loop that maintains the stable outgrowth of the limb bud. In agreement with previous reports (Glotzer et al., 2022; Lovely et al., 2022; Suzuki et al., 2024), *Fgf8* expression was observed in the anterior region of the limb mesenchyme, whereas *Fgf10* expression was detected uniformly throughout the mesenchyme in salamander hindlimb bud. We also examined the expression patterns of *Hoxa11* and *Hoxa13*, two key genes involved in proximal-distal axis patterning during limb development (Figure 4E–G). Generally, *Hoxa11* is associated with zeugopod formation, whereas *Hoxa13* is involved in vertebrate autopod development (Davis et al., 1995; Fromental-Ramain et al., 1996). We found that *Hoxa11* expression was distributed in the intermediate region of the newt hindlimb bud, whereas *Hoxa13* expression was localized to the distal region, consistent with a previous report (Takeuchi et al., 2022). Furthermore, we simultaneously detected both *Shh* and *Fgf8* signals in the same tissue section (Figure 4H–J). In salamanders, *Shh*, which plays a crucial role in anterior-posterior (AP) axis specification during limb development, is expressed in the posterior mesenchyme (Glotzer et al., 2022; Lovely et al., 2022). As previously reported, we observed that *Shh* transcripts were localized to the posterior mesenchyme, while the *Fgf8* signals were detected in the anterior mesenchyme of the newt hindlimb bud.

**Figure 4.**
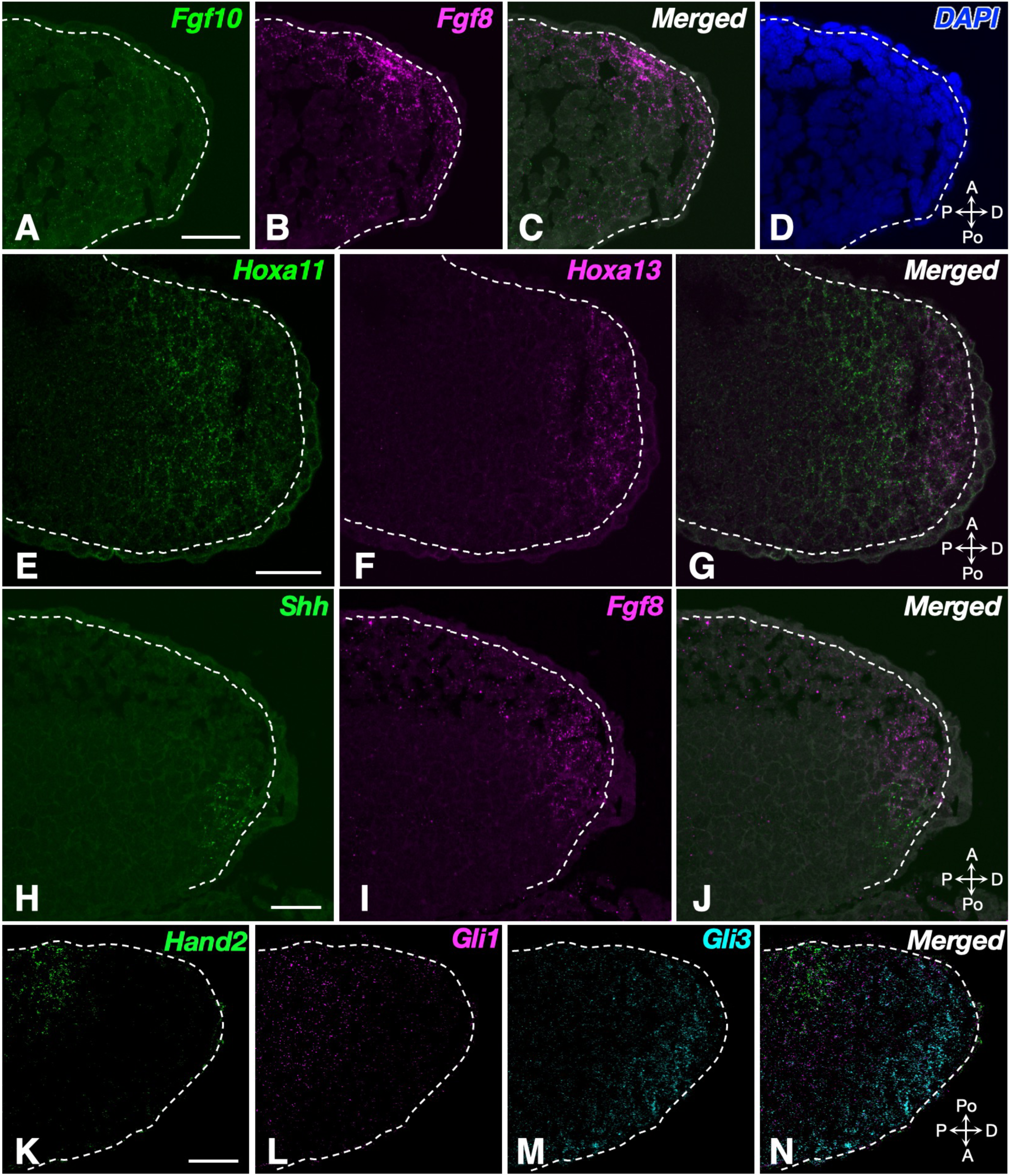
mRNA detection of limb development-related genes by a safer *in situ* HCR protocol in newt hindlimb bud sections. (A–C) Dual detection of *Fgf8* (magenta, B) and *Fgf10* (green, A) in newt hindlimb bud. *Fgf8* is enriched in the anterior mesenchyme, whereas *Fgf10* is detected sparsely throughout the bud; the merged image is shown in C. (D) Nuclear counterstaining with DAPI. (E–G) Complementary expression domains of *Hoxa11* (green) and *Hoxa13* (magenta) along the proximal-distal axis; the merged image is shown in G. (H–J) Mutually exclusive expression of *Shh* (green) and *Fgf8* (magenta) in the posterior and anterior mesenchyme, respectively; the merged image is shown in J. Sections in A–D and H–J were post-fixed with ALTFiX, whereas sections in E–G were fixed with PAXgene®. (K–N) Triple-plexed visualization of SHH-pathway transcripts by a safer *in situ* HCR protocol in newt hindlimb bud sections post-fixed with ALTFiX. (K–M) Individual channels showing *Hand2* (green, K), *Gli1* (magenta, L), and *Gli3* (cyan, M) mRNAs. (N) A merged view of all three channels. *Hand2* labels the posterior putative *Shh*-expressing domain, while *Gli1* and *Gli3* are positively and negatively regulated by SHH signaling, respectively, resulting in sharply segregated expression domains along the anterior-posterior axis. Dotted white lines indicate the epithelial-mesenchymal boundary. Abbreviations: A, Anterior; Po, Posterior; P, Proximal; D, Distal. Scale bars: 50 µm.

We further performed simultaneous triple-plex detection of *Hand2*, *Gli1*, and *Gli3* in the newt hindlimb bud. *Hand2* and *Gli1* are known to be expressed in the posterior mesenchyme under the control of SHH signaling, whereas *Gli3* is activated in the anterior mesenchyme (Ahn & Joyner, 2004; Galli et al., 2010). Therefore, these genes are generally employed as readouts of SHH signaling in limb development. Consistent with these previous reports, our results confirmed these expression patterns in the newt hindlimb (Figure 4K–N). *Hand2* exhibited a relatively localized signal in the posterior region (Figure 4K), whereas *Gli1* showed a gradient extending from the posterior toward the anterior region (Figure 4L). In contrast, *Gli3* was detected in the anterior region opposite to the *Shh*-expressing region, consistent with its negative regulation by SHH signaling (Figure 4M, N). Even when using the triple-plex fluorophores Alexa Fluor 488/514/546, which have overlapping excitation and emission spectra, the *in situ* HCR images clearly delineated the three distinct gene expression domains and no significant fluorescence bleed-through was observed.

### 2.5 Application of the safer protocol to SABER-FISH mRNA detection in model organisms

To demonstrate the utility of our safer FISH protocol, we examined its applicability to SABER-FISH. A key feature of SABER-FISH is “branching” in which branched secondary probes hybridize to multiple fluorophore-labeled detection probes, thereby greatly enhancing signal intensity. In this study, we performed one round of branching for all SABER-FISH experiments to detect mRNA signals. To validate this application, we employed a pre-fixed frozen section preparation protocol in the medaka retina, an experimental system in which the success of SABER-FISH has been previously established (Fukuda et al., 2025). In this experiment, the 4% PFA typically used in the routine pre-fixation protocol for the medaka retina was simply replaced with ALTFiX. We targeted retinal cell-type-specific marker genes, including *gja10b* for horizontal cells and *cabp5a* for bipolar cells (Figure 5A–C) (Glasauer & Neuhauss, 2016; Klaassen et al., 2016). As expected, in ALTFiX-fixed retinas, *gja10b* was detected at the boundary of the outer plexiform layer (OPL) and the inner nuclear layer (INL), and *cabp5a* was detected in the outer part of the INL (Figure 5A–C). Notably, autofluorescence from the photoreceptor inner segments was easily distinguishable from the punctate FISH signals. These results demonstrate that the pre-fixed frozen section protocol using a safer fixative is compatible with SABER-FISH in the medaka retina. Importantly, no special modifications were required when replacing the toxic 4% PFA with the safer ALTFiX in the medaka tissue preparation protocol.

**Figure 5.**
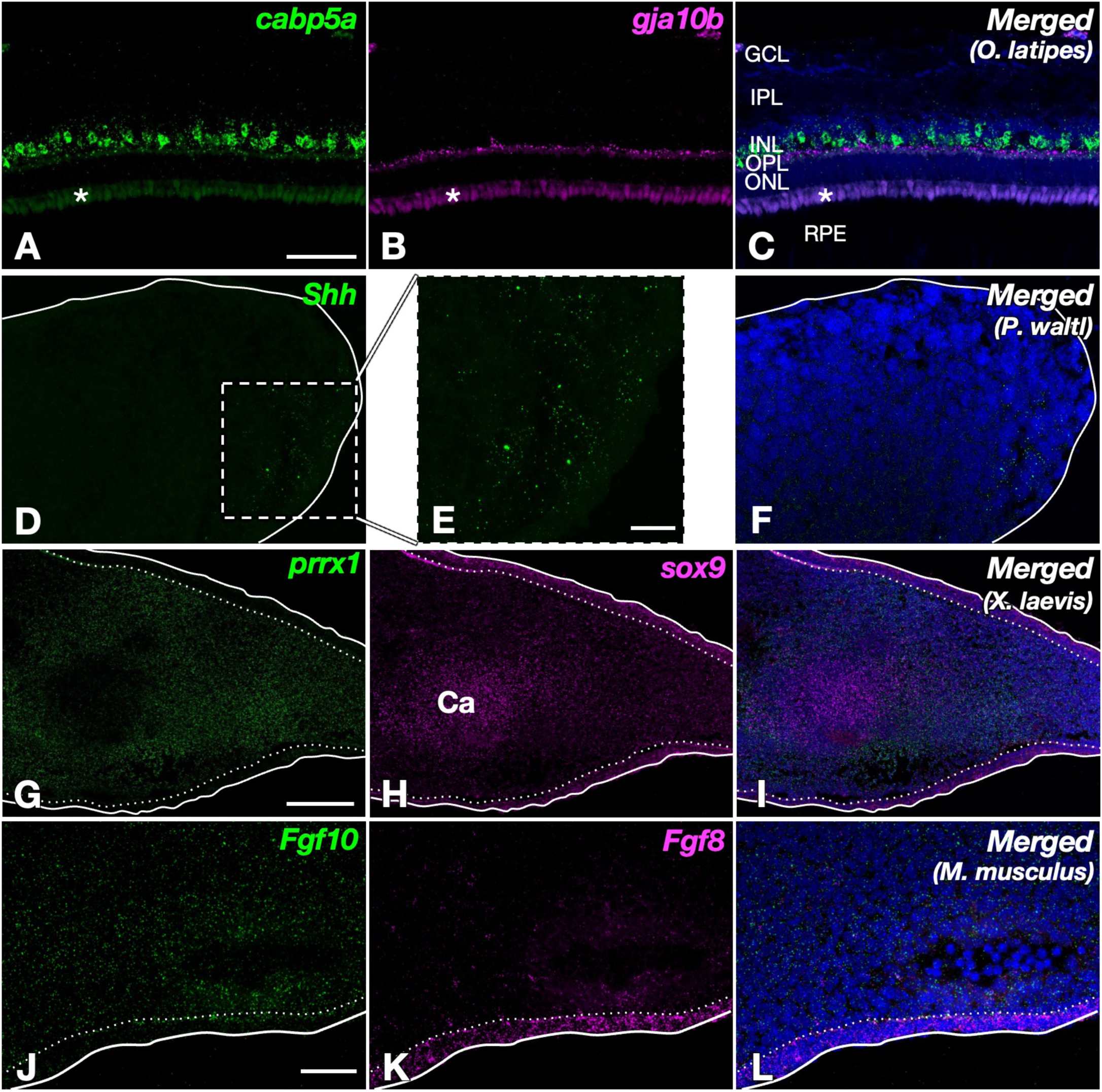
Versatile mRNA detection in diverse vertebrate species using the optimized SABER-FISH protocol. Tissue sections were pre- or post-fixed with ALTFiX. (A–C) Detection of *cabp5a* (green, A) and *gja10b* (magenta, B) in the medaka (*O. latipes*) retina. (C) Merged image with DAPI. Asterisks indicate autofluorescence from the inner segments of photoreceptor cells. Abbreviations: RPE, retinal pigment epithelium; ONL, outer nuclear layer; OPL, outer plexiform layer; INL, inner nuclear layer; IPL, inner plexiform layer; GCL, ganglion cell layer. (D–F) *Shh* detection in the posterior region of stage 40 newt hindlimb bud. (E) Magnified view of the area boxed in (D). (F) Merged image with DAPI. (G–I) Simultaneous detection of *prrx1* (green, G) and *sox9* (magenta, H) in stage 53 frog hindlimb bud. *prrx1* is expressed in mesenchymal cells, whereas *sox9* is localized to chondrocytes. (I) Merged image with DAPI showing expression patterns along the dorsal–ventral axis. Abbreviations: Ca, cartilage. (J–L) Detection of *Fgf10* (green, J) and *Fgf8* (magenta, K) in mouse (*M. musculus*) limb buds. (L) Merged image with DAPI. Solid white lines outline the limb buds, and dotted white lines indicate the epithelial–mesenchymal boundary (D–L). Scale bars: (A, G, J) 50 µm; (E) 20 µm.

Next, to demonstrate the broader utility of our safer FISH protocol across other vertebrates, we evaluated its applicability to the SABER-FISH in amphibians. First, to assess the effectiveness of this approach, we examined *Shh* expression in stage 40 newt hindlimb bud. SABER-FISH clearly detected *Shh* signals in the limb buds (Figure 5D–F), showing the same distribution pattern as shown in Figure 4H. Next, we investigated the expression patterns of *prrx1* and *sox9* in stage 53 frog hindlimb bud (Figure 5G–I). During frog limb development, *prrx1* is broadly expressed in the limb mesenchyme, whereas *sox9* is exclusively expressed in aggregating chondrocytes (Satoh et al., 2006; Suzuki et al., 2007). As expected, SABER-FISH accurately recapitulated these distinct expression patterns of each gene in the developing amphibian hindlimb.

To further confirm the broad applicability of our protocol to mammals, we applied SABER-FISH to mouse limb sections targeting *Fgf10* and *Fgf8*. These genes are essential regulators of limb development, with *Fgf10* expressed in the mesenchyme and *Fgf8* restricted to the apical ectodermal ridge (AER) (Crossley & Martin, 1995; Lewandoski et al., 2000; Min et al., 1998). Previous studies have demonstrated *Fgf10* detection in PFA-fixed mouse lacrimal gland samples, and we similarly detected signals in limb sections using our safer protocol with a probe targeting the same sequence (Ikeda et al., 2026). As expected, SABER-FISH using our safer protocol clearly captured these distinct expression patterns for both *Fgf10* and *Fgf8* (Figure 5J–L). Notably, as observed in medaka and amphibians, our protocol was readily adaptable to the mouse tissue preparation without the need for specialized modifications.

### 2.6 Comprehensive multiplexed gene detection via the integration of *in situ*

#### HCR and SABER-FISH using the safer protocol

To evaluate whether the safer protocol enables the seamless integration of *in situ* HCR and SABER-FISH to achieve the comprehensive multiplexed gene detection, we performed the combined detection of two methods on the same tissue sections (SABER-HCR). Limb bud sections from E12.5 mouse embryos were analyzed (Figure 6). *Sox9* and *Col2a1* transcripts detected by *in situ* HCR were largely co-expressed in chondrogenic domains (Figure 6A), which is consistent with the results shown in Figure 2D–F. Although *Gdf5* plays a critical role in joint formation and is restrictedly expressed in prospective joint interzones (Storm et al., 1994; Storm & Kingsley, 1996), its expression pattern detected by SABER-FISH exhibited a pattern surrounding the developing digits at this stage (Figure 6B, C). Importantly, signals obtained by SABER-FISH and *in situ* HCR were clearly distinguishable within the same tissue sections. No apparent signal interference or cross-reactivity between the two amplification systems was observed.

**Figure 6.**
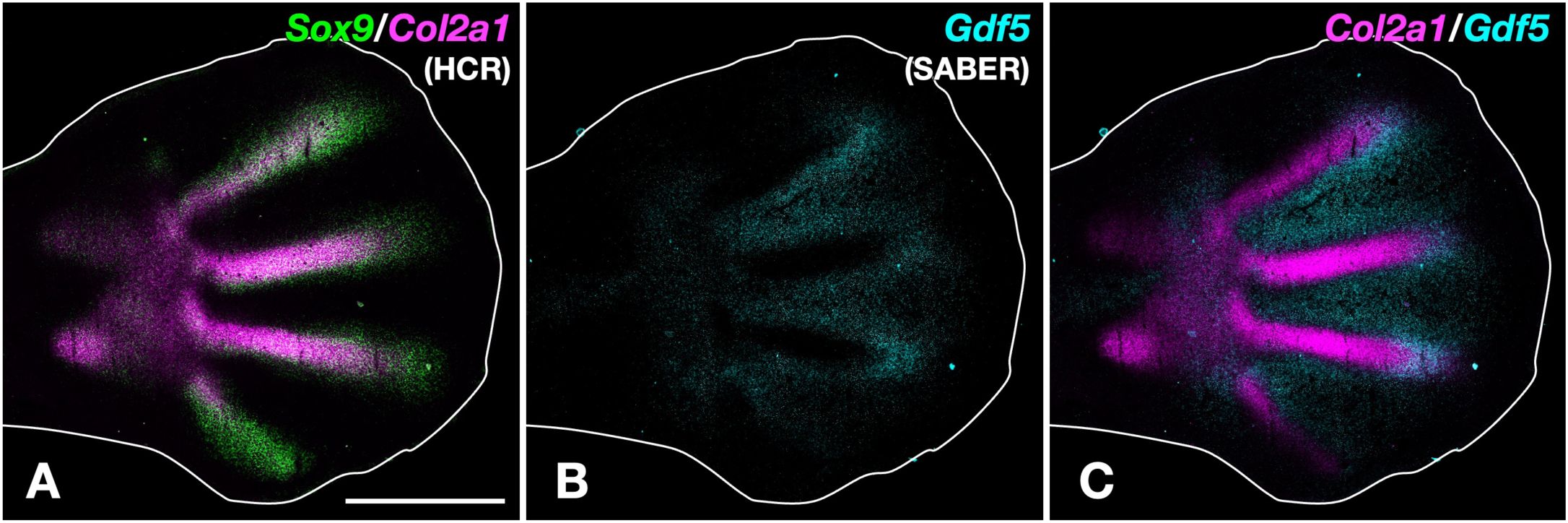
Comprehensive multiplexed gene detection of *in situ* HCR and SABER-FISH signals in E12.5 mouse limb buds (SABER-HCR). (A) Detection of *Sox9* (green) and *Col2a1* (magenta) transcripts by *in situ* HCR. (B) Detection of *Gdf5* transcripts by SABER-FISH. (C) Merged image of *Col2a1* (HCR signal; magenta) and *Gdf5* (SABER signal; cyan. Solid white lines outline the limb bud. Scale bar: 500 µm.

## 3. Discussion

mRNA FISH has been an essential research strategy in current biology, enabling the analysis of tissues, organs, and even whole organisms at single-cell resolution while simultaneously permitting examination of the expression of a limited set of genes in the same sample. *In situ* HCR, RNAscope, and SABER-FISH, each of which employs various amplification techniques, have been widely applied in various biological research fields (Choi et al., 2018; Kishi et al., 2019; Wang et al., 2012). The simultaneous analysis of 2–4 genes in a single observation, although limited by the combination of fluorophores and their filters, is becoming increasingly routine. Given these developments, improved FISH protocols are expected to continue to grow in importance as research tools. The low-toxicity FISH protocol developed in this study offers a beneficial option for safeguarding the health of researchers performing routine experiments.

Glutaraldehyde and paraformaldehyde (formalin) are among the most widely used fixatives in various histological techniques, including FISH, due to their effectiveness in preserving tissue. In developmental biology, it has been reported that PFA provides superior signal detection in HCR compared to other fixatives (Echeverria et al., 2025). However, their high toxicity and volatility necessitate careful handling to ensure the safety of researchers. To address this issue, we developed a safer protocol by replacing a toxic fixative with less toxic alternatives. One major concern when switching to alternative fixatives was the potential reduction in detection sensitivity for *in situ* HCR and SABER-FISH. Nonetheless, our protocol successfully detected the expression of even low-abundance transcripts (e.g., morphogens and transcription factors) in tissue sections, which are typically difficult to identify using RNA-seq analysis. Despite being an aldehyde-based compound, glyoxal is a relatively low-toxicity and robust cross-linking fixative that has recently gained attention in the life sciences (Konno et al., 2023; Richter et al., 2018). Glyoxal not only serves as an effective substitute for formaldehyde (formalin) in mRNA FISH (Amin et al., 2023), but it has also been reported to enhance signal sensitivity in certain cases (Yao et al., 2021). Moreover, excessive fixation with strong cross-linking agents such as formalin often causes signal attenuation in ISH. To mitigate this issue, tissues fixed with formalin generally require additional steps to partially reverse cross-linking, thereby improving probe accessibility to mRNA. These additional steps commonly involve proteolytic enzymes such as proteinase K, pepsin, and trypsin, which are widely employed in ISH. However, enzymatic treatments damage tissues and require careful optimization of parameters such as treatment time, concentration, and temperature, depending on the sample and the individual target genes. In contrast, our protocol using safer fixatives eliminates the need for proteolytic treatment using the enzymes. By avoiding enzymatic digestion, the protocol not only simplifies the experimental workflow but also provides the additional advantage of better preservation of tissue morphology. Importantly, our protocol is highly compatible with integrated mRNA FISH and IHC, facilitating the phenotypic characterization of EGFP-labeled cell populations in transgenic models. This versatility also extends to the spatial validation of scRNA-seq results, enabling the multimodal detection of target transcripts (mRNA) and specific cell-surface antigens (protein) at single-cell resolution *in situ*.

Methanol is a highly toxic organic solvent that poses significant health risks in the laboratory. Nevertheless, conventional FISH protocols require methanol treatment for fixation, permeabilization, delipidation, and dehydration, because methanol effectively increases the probe penetration and enhances the signals. Indeed, in zebrafish whole-mount FISH, methanol treatment is critical (Oka & Sato, 2015). On the other hand, because it uses glyoxal-based fixation, our protocol does not require methanol treatment.

In this study, we evaluated the application of our low-toxicity protocol to both *in situ* HCR and SABER-FISH using frozen tissue sections across various model animals. *In situ* HCR is a user-friendly method because the design and synthesis of the probes (initiator probe and amplifier) can be outsourced to suppliers or designed independently by researchers, enabling us to readily perform this technique. In various model animals widely used in developmental biology, including not only *M. musculus* but also *X. laevis*, *A. mexicanum*, *P. waltl*, and *O. latipes*, *in situ* HCR technology enables the detection of mRNA expression with high sensitivity and resolution (Aztekin et al., 2021; Glotzer et al., 2022; Jaeger et al., 2025; Lee et al., 2023; Lovely et al., 2022, 2023). Moreover, an improved version of *in situ* HCR that enhances probe permeability by using shortened hairpin amplifier probes was recently reported (Tsuneoka & Funato, 2020). Integrating this improved *in situ* HCR with our protocol could not only enhance stability but also improve the consistency and reliability of signal detection, thereby increasing the success rate of experiments. On the other hand, SABER-FISH is somewhat less user-friendly, as it requires researchers to design and synthesize probes themselves by using *in vitro* primer exchange reactions (PER) (Kishi et al., 2018). Nevertheless, the PER concatemers, branch probes, and fluorescent imager probes can be used following the procedures described in the original publication (Kishi et al., 2018), and the required oligos can be synthesized by ordering sequences from a supplier. One key advantage of SABER-FISH is that the use of branch probes allows an increase in the number of hybridized fluorescent imager probes. This can amplify signals and enable the detection of low-abundance mRNAs that are challenging to identify with *in situ* HCR. However, when we designed probes for the same gene ORF and used them to compare the performance of *in situ* HCR and SABER-FISH, we observed instances where only one method successfully detected the target (data not shown). Although the exact reason for this differential efficacy remains unknown, it is possible that the position or number of probes designed by each method affects the efficiency of hybridization against certain target mRNAs. The present study demonstrates that *in situ* HCR and SABER-FISH can be applied simultaneously within the same tissue sections using the safer protocol, without apparent signal interference. The two amplification systems produced clearly distinguishable signals that were consistent with known spatial expression patterns across multiple vertebrate species, supporting their compatibility. These findings suggest that *in situ* HCR and SABER-FISH are not fully interchangeable but rather complementary techniques. Therefore, the ability to use both methods as alternative options provides a practical advantage, increasing the likelihood of successful detection across a broad range of target genes.

## 4. Limitations of the study

Certain limitations of our low-toxicity FISH protocol should be noted. First, the protocol has not yet been validated for use with whole-mount or paraffin-embedded samples in vertebrates, and further investigation of these applications is warranted. Nevertheless, glyoxal-based fixation has proven effective in whole-mount FISH (WISH) of *Drosophila* embryos (Amin et al., 2023), suggesting that it may also be applicable to WISH on vertebrate embryos and tissues. Second, particularly for soft and fragile materials such as embryos, it may be preferable to fix the samples before embedding them. Third, despite the fact that we observed virtually no loss of FISH signal sensitivity in our experiments, glyoxal-based fixation is slightly less effective at preserving cellular and tissue morphology, particularly nuclear structure, compared to paraformaldehyde or other aldehyde fixatives. Thus, the advantages and disadvantages are essentially a trade-off: the modest compromise in morphology is offset by the ability to conduct research while markedly reducing researcher exposure to toxic formalin and methanol. Fourth, a safety concern remains regarding the continued use of formamide, an organic solvent widely employed to facilitate probe hybridization to target genomic DNA and RNA in conventional FISH protocols. High concentrations of formamide can also pose health risks to researchers and lab staff conducting experiments. Consequently, several ISH protocols that utilize alternative organic solvents or reagents such as urea and ethylene carbonate have been described (Matthiesen & Hansen, 2012; Sinigaglia et al., 2018). Notably, ethylene carbonate both promotes hybridization and shortens hybridization times in *in situ* HCR (Mikami et al., 2025). Future efforts will focus on incorporating this reagent into our FISH protocol to further improve its safety and ease of use in the laboratory.

## 5. Experimental Procedures

### 5.1 Animals

Iberian ribbed newts (*Pleurodeles waltl*) were provided by the National BioResource Project (NBRP), Amphibian Research Center, Hiroshima University. African clawed frog (*Xenopus laevis*) tadpoles and adults were maintained in-house, while knock-in tadpoles (*sox2:egfp*) were generated at the University of Hyogo (Mochii et al., 2024). Medaka (*Oryzias latipes*, strain d-rR) and mice (*Mus musculus,* strain C57BL/6) were maintained and bred at Okayama University. For deep anesthesia of amphibians and medaka, 0.2% MS-222 (Sigma, USA) and 0.1% (v/v) 2-phenoxyethanol were used. For mice, pregnant C57BL/6 wild-type females were euthanized by cervical dislocation to collect E12.5 embryos. Developmental stages were determined according to established staging tables for *P. waltl* and *X. laevis* (Nieuwkoop & Faber, 1994; Shi & Boucaut, 1995). Animals were euthanized according to methods approved by the Institutional Animal Care and Use Committee, followed by the sampling of tissues and organs. All animal care was approved by the Institutional Animal Care and Use Committee of the National Institutes of Natural Sciences, University of Hyogo, and Okayama University.

### 5.2 Preparation of cryosections

All tissues for post-fixation were embedded in Tissue-Tek O.C.T. compound (Sakura Finetek, Japan) and immediately frozen on an aluminum plate cooled with liquid nitrogen. Frozen amphibian limb buds were sectioned at a thickness of 14 µm using a Cryostar NX70 cryostat (Thermo Fisher Scientific, USA) and mounted on MAS-coated or CREST-coated glass slides (Matsunami Glass, Japan). Frozen mouse limb buds were sectioned at a thickness of 10 µm using a cryostat (Polar; Sakura Finetek, Japan) and mounted on CREST-coated glass slides. Mounted sections were fixed at room temperature for 15 min using one of two formaldehyde-free tissue fixatives, ALTFiX (FALMA, Japan) or PAXgene® (Qiagen, Germany). 4% paraformaldehyde was also used for comparison in Figure 2. For tadpole samples, pre-fixation was performed prior to dissection by immersing the animals in ALTFiX overnight at 4°C. After pre-fixation, tissues were dissected, embedded in O.C.T. compound, and frozen at −80°C prior to cryosectioning. For medaka retina samples, dissected tissues were pre-fixed by immersion in ALTFiX overnight at 4°C. Following pre-fixation, retinas were cryoprotected by immersion in 20% sucrose in phosphate-buffered saline (PBS) overnight, embedded in O.C.T. compound, and frozen at −80°C prior to cryosectioning. Frozen medaka retinas were sectioned at a thickness of 15 µm using a cryostat (Polar; Sakura Finetek, Japan) and mounted on CREST-coated glass slides. The tissue sections were dehydrated through a graded ethanol series (50%, 70%, 100% and 100%). After air-drying for 30 min, the slides were outlined with an ImmEdge Hydrophobic Barrier PAP Pen (Vector Laboratories, USA). Sections were then immersed in a modified detergent solution (Table S1) (Bruce & Patel, 2022; Morabito et al., 2023) for 30 min at room temperature and washed twice with PBS.

### 5.3 Probe design

For SABER-FISH probe design, probes targeting *prrx1* and *sox9* of *X. laevis*, *Shh* of *P. waltl*, *gja10b* and *cabp5a* of *O. latipes* were designed using the blockParse script from the OligoMiner package (Beliveau et al., 2018). Probes targeting *Fgf8* and *Gdf5* of *M. musculus* were designed using the PaintSHOP web tool (Hershberg et al., 2021). The probe sequence targeting *Fgf10* of *M. musculus* was designed based on previously reported sequences (Ikeda et al., 2026). All coding sequences (CDS) of the target genes were retrieved from Xenbase (Fisher et al., 2023), the two *P. waltl* genome projects (Brown et al., 2025; Kimura et al., 2025), and the NCBI nucleotide database.

### 5.4 *in situ* Hybridization chain reaction (*in situ* HCR)

This protocol is based on a protocol developed by Molecular Instrument (USA), and HCR probes, probe hybridization buffer, and amplification buffer were obtained from Molecular Instrument. The permeabilized samples were pre-hybridized with a probe hybridization buffer for 10 min at 37°C. A probe solution (comprising 0.4 pmol of each initiator probe in 100 μL of probe hybridization buffer) was applied, and the samples were incubated overnight at 37°C. Following hybridization, the slides were washed three times in a pre-warmed wash solution (10% formamide, 2× saline-sodium citrate [SSC], and 0.1% Tween 20) for 1 h each at 37°C. Sections were then incubated in 5× SSC containing 0.1% Tween 20 (SSCT) for 30 min at 37°C, followed by an additional wash in 5× SSCT for 5 min. The samples were incubated with an amplification buffer at room temperature for 1 h. After heat denaturation of hairpin probes at 95°C for 90 seconds, 100 µL of hairpin solution (containing 2 µL each of 3 µM hairpin h1 and h2 probes in 100 µL of amplification buffer) was applied, and the samples were incubated overnight at room temperature in the dark. For fluorescence detection, hairpin probes labeled with Alexa Fluor dyes (Alexa Fluor 488, 514, 546, and 647; Molecular Instruments) were used for HCR signal amplification. After amplification, sections were washed twice in 5× SSCT for 30 min and once for 5 min at room temperature. For tadpole brain sections, IHC was performed following these washes. The sections were incubated for 1 h at room temperature with an anti-GFP mAb-Alexa Fluor™ 647 (1:500; MBL Co., Ltd., Japan) diluted in Can Get Signal Immunostain Solution B (TOYOBO Co., Ltd., Japan). After antibody incubation, sections were washed three times in PBS containing 0.1% Tween 20 (PBST). Finally, the slides were mounted using ProLong™ Gold Antifade Mountant with DNA Stain DAPI (Thermo Fisher Scientific, USA).

### 5.5 Signal Amplification By Exchange Reaction (SABER)-FISH

Primer exchange reactions were performed following previous reports (Ikeda et al., 2026; Fukuda et al., 2025; Kishi et al., 2019). Fluorescent signals were amplified and detected with one branching round as follows. Sections were treated with the detergent solution at room temperature, followed by three rinses in PBST. They were pre-hybridized in a pre-hybridization buffer (Prehyb; 40% formamide, 2× SSC, 1% Tween 20, 1× Denhardt’s solution, 100 μg/mL heparin sodium, and 0.5 μM random 20-mer oligo DNA) at 43°C for 10 min, then incubated with the respective probes at 1 μg/mL in a primary probe hybridization buffer (Hyb1; Prehyb containing 10% dextran sulfate [FUJIFILM Wako Pure Chemical Corporation 197-09984]) at 43°C overnight. Post-hybridization washes were performed in a wash hybridization buffer (Whyb; 40% formamide, 2× SSC, 1% Tween 20) and 2× SSCT. For signal amplification, branch oligos were applied at 37°C overnight, followed by SSCT washes. For experiments combining SABER-FISH and *in situ* HCR (SABER-HCR; Fig. 6), branch oligos (1 μg/mL in Hyb1) were applied together with HCR initiator probes under the same hybridization conditions at 37°C overnight. After washing with Whyb and SSCT and rinsing with PBST, fluorophore-conjugated oligos (Imager oligos, 0.1 μM) diluted in a secondary hybridization buffer (Hyb2; 0.1% Tween 20, 1× Denhardt’s solution, 100 μg/mL heparin sodium, and 0.5 μM random 20-mer oligo DNA in PBS) were added for 45 min at 37 °C to detect SABER signals. Sections were washed with PBST, after which HCR amplification hairpins were applied and incubated overnight at room temperature. Finally, the slides were mounted using ProLong™ Gold Antifade Mountant with DNA Stain DAPI. For medaka retina sections, however, the tissues were counterstained with DAPI (0.2 µg/mL), rinsed in PBST, and coverslipped with a homemade polyvinyl alcohol/glycerol mounting medium. The oligonucleotide sequences for the SABER-FISH probes are described in Table S2.

### 5.6 Microscopy and image acquisition

Confocal imaging was performed using two laser scanning confocal microscope systems: a Nikon A1R (Nikon, Japan) and a Leica STELLARIS8 (Leica Microsystems, Germany). The two systems were used depending on the combinations and number of fluorophores to be imaged. The Nikon A1R was equipped with 20× dry and 40× water-immersion objectives and laser lines at 405, 457, 488, 514, 561, and 640 nm. This system was primarily used for *in situ* HCR and SABER-FISH experiments with up to three fluorophores (Alexa Fluor 488, 546, and 647), together with DAPI nuclear staining. The Leica STELLARIS8 was equipped with a HC PL APO 40×/1.30 oil-immersion CS2 objective (working distance 0.17 mm), and used a 405 nm laser and a white light laser (440–790 nm). This system was used for experiments requiring four-color HCR or combined three-color HCR and one-color IHC, together with DAPI (Alexa Fluor 488, 514, 546, and 647). Images acquired on the STELLARIS8 were collected and processed using LAS X software (Leica Microsystems, Germany). For low-magnification overview images, tile scanning was performed and individual tiles were stitched using the manufacturer’s software (NIS-Elements for Nikon A1R and LAS X for STELLARIS8). For combined *in situ* HCR and IHC experiments, EGFP fluorescence, HCR signals, and IHC signals were sequentially acquired from the same tissue sections. For comparisons between different fixation conditions (PFA and ALTFiX) or detection methods, images were acquired using identical acquisition parameters within each imaging system. Image processing was performed using Fiji (Schindelin et al., 2012). Only linear adjustments of brightness and contrast were applied equally to entire images.

## CRediT authorship contribution statement

Conceptualization: KTS, Methodology: KS, KTS, Experiments: AC, RM, NK, AT, AO, MS, YS, KS, KTS, Visualization: AC, RM, NK, KS, KTS, Funding acquisition: KS, KTS, Project administration: KS, KTS, Supervision: MM, HO, Writing – original draft: AC, RM, KS, KTS, Writing – review & editing: All authors

## Acknowledgments

*P. waltl* newts were provided by the Hiroshima University Amphibian Research Center, and the National BioResource Project of MEXT. This research was supported by the following funding sources: JST, CREST Grant Number JPMJCR2025 to K.T.S. and JSPS KAKENHI Grant Numbers 21H03829 and 25K02288 to K.T.S. and 23K05850 to K.S.; the NIBB Collaborative Research Program (25NIBB338) to M.M.; Joint Research of the Exploratory Research Center on Life and Living Systems (ExCELLS) Grant Numbers 23-S6 and 22-S3 to K.T.S.; Frontier Photonic Sciences Project of National Institutes of Natural Sciences (Grant Number 01212503) to K.S.; a grant from the Japan Foundation for Applied Enzymology to K.S.; and a grant from the Nagahisa Science Foundation to A.C. We thank Dr. Yoshihiro Morishita and Ms. Kaori Niimi (RIKEN BDR) for technical advice of cryosection and KN international (Japan) for their professional assistance with basic English proofreading. Confocal images were acquired at the Optics and Imaging Facility of the NIBB Trans-Scale Biology Center.

## Declaration of generative AI and AI-assisted technologies

The authors acknowledge the use of ChatGPT and Gemini for language editing support. All AI-assisted texts were reviewed and revised by the authors. The authors take full responsibility for the contents.

---

**Supplementary figure 1.** Combined *in situ* HCR and immunohistochemistry in brain sections from stage 50 wild-type frog tadpole. (A–G) *In situ* HCR for *sox2* and *shh* combined with anti-EGFP immunohistochemistry in a brain section. (A, D, and F) Low-magnification fluorescence images of a brain section. (B, C, and E) Enlarged views of the region outlined by a dashed line in A, D, and F. (A, B) DAPI nuclear staining. (D, E) EGFP channel. (C) *In situ* HCR signals for *sox2* (magenta) and *shh* (cyan) mRNAs. (F, G) Immunofluorescence using an anti-EGFP antibody; no signal was detected. Scale bars: (A, D, and F) 200 µm, (B, C, E, and G) 50 µm.

