## Supplementary material for "A safer fluorescent *in situ* hybridization protocol for cryosections": Sa

### Figure S1

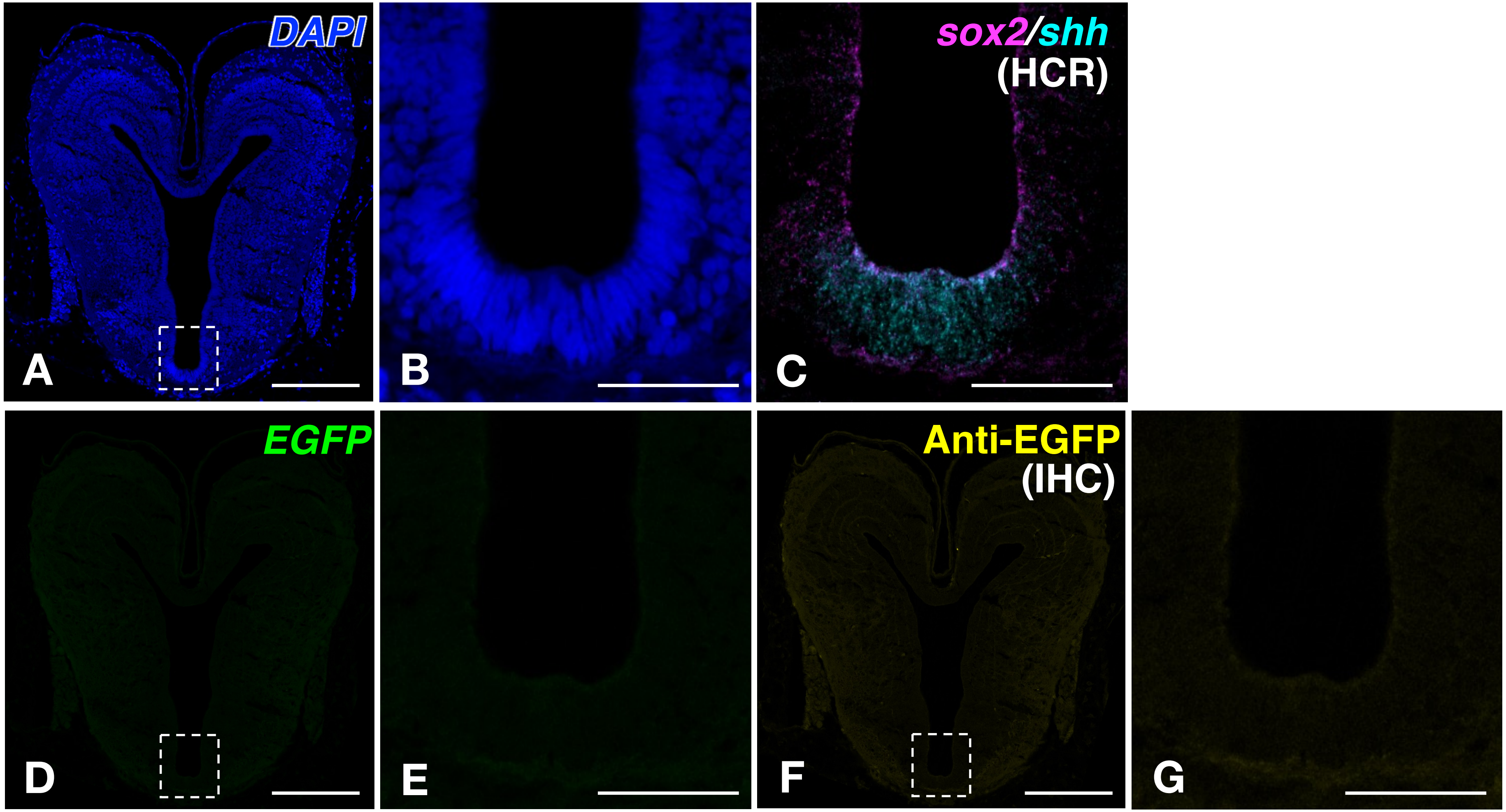

### Supplementary Tables

**Table S1.** Composition of detergent solution (up to 1 L with sterile distilled water)

| Reagents | Concentration (Final) |
|---|---|
| SDS | 0.1 % |
| Tween 20 | 0.5 % |
| Tris-HCl (pH7.5) | 50 mM |
| EDTA (pH8.0) | 1 mM |
| NaCl | 150 mM |

**Table S2.** Oligonucleotide sequences for SABER-FISH probes

| Animal | Gene | Sequence |
|---|---|---|
| *Oryzias latipes* | *cabp5a* | 5'-TTAAAGTGCAGAGACTCAGACTGTCCTGCCCGCTGC-3' |
| | | 5'-AATCAGAGCCGCCTCTCTTCAGCCAAGACAGCTTGG-3' |
| | | 5'-GTCCTCCTTTGTCCTTCTGTTTCTTGCTCTTCCTCTTAG-3' |
| | | 5'-AACCCTGCAGTTCAGATTCAGCAGCTGACTCCTGCG-3' |
| | | 5'-GTGATTTCTCCTCATCAGTCAAGAGCGTCACACGTG-3' |
| | | 5'-AAATGCACGCTGCACCGAAACTCATTTCTTCTTCTG-3' |
| | | 5'-TCATCAATCTCATCGTCAGCCAACTCTCTGGAAATGTTTT-3' |
| | | 5'-TTGTCTTTGTCAAACTCATTGAAAGCGTCCCGCAGC-3' |
| | | 5'-AGATTCCCCAGATCCTTGCAGGTGATCAGCCCATCC-3' |
| | | 5'-TCCATCTCCGTGGGCATGTAACCCATGGTCCTCATC-3' |

| | | |
|---|---|---|
| | | 5'-AGGTTCATGTTGATGTTCTGGCTCAGTTCGATCAGC-3' |
| | | 5'-AGTTCCACAAAGTCCTCAAAGTCAACTCTGCCACCA-3' |
| | | 5'-TTGAGCTCCTTCACGCCAATCATCCCAGCTGTTTCCG-3' |
| | | 5'-CCCATCTCCATCCATGTCAAACTCTTTGAAGGCATCT-3' |
| | | 5'-GTTCATGGCTGCTCGCAGCTCTTCCGTTGTGATTTC-3' |
| | | 5'-GATCTCTCTGCGGCTCATGTGCTCTCCCATCAGCTT-3' |
| | | 5'-GTCTCCATTGTCATCGGCTTCTTTCACAATGGCGTC-3' |
| | | 5'-CCATCACTGACGAGACATCATTCGGACAAATTCTTCAAAGT-3' |
| | | 5'-GCAAATAAGCATTCTTGAAAGGTGCTGTTTCCGCGT-3' |
| *Oryzias latipes* | *gja10b* | 5'-CTCCTCTAAGATACTTCCCAGAAGGTTCCAGTCCCC-3' |
| | | 5'-GCCAGATCTTCCCCACAATGGTGGAATGAATATGGAC-3' |
| | | 5'-CCCAATACCAGCATCCGGAAAATGAAGAGGATGGTAA-3' |
| | | 5'-CAAACTCACTCTGCTCGTCATCCCAGACGTCCTCGG-3' |
| | | 5'-GAGATGGGAAACGCCTGGTCGTAGCAGACGGTCTTG-3' |
| | | 5'-ACGAAGATGATCTGAAGGACCCAGTACCGGATGAGG-3' |
| | | 5'-TCTCTCCTTTTCCAGGGTCCTCAGGCGGTACAGAGC-3' |
| | | 5'-CTCCAGCTCAGCTTTCAGGCAGGCTTTCTTCCTGTG-3' |
| | | 5'-TCTCGCTCGATCCTCTTGTGGTCCTCCTGGTCTGTC-3' |
| | | 5'-TTTTGGACTTTCTTCTGCTCGTCCAGCTTTCTGAGT-3' |
| | | 5'-AGACATAAGTGCGCAACAAGGAGCCCCTCAGAGGAG-3' |
| | | 5'-AGCCAACCTCCACCACAGATCTGGTCAGGATATGGA-3' |
| | | 5'-GTCCAATGCCATACAGAGCACACTGACCAATGATGA-3' |
| | | 5'-GACAAGGCAAGCGTTCACACTTGTACATGGAGGACA-3' |
| | | 5'-CTGTCGGTCGCGACACAAAACAGTCCACAACGTTTG-3' |

| | | |
|---|---|---|
| | | 5'-AGCTATGACCAGCATGAAGACCATGAAAACGTTCTTCT-3' |
| | | 5'-GGTGGAAGATCTCCATAATGTTGAGGAACAGAGAAACTCC-3' |
| | | 5'-GTATCCGTACAAGATGTCCTTGATCCGCTTCACCCC-3' |
| | | 5'-TCTTCTTGGACCTGTACACGCTATCTTCGTCTCCATATTT-3' |
| | | 5'-GAGCAGCTCATCTGAGTGAGCTGCATCAGCCTCTGG-3' |
| | | 5'-GACCAAATGCAAAGGCATGGTCTCCCCGTGACTGTG-3' |
| | | 5'-TGGGCTGTGATGAAATCCTCTTGGTTTTGAGCGGGG-3' |
| | | 5'-TGGGTGTCAGTTTGGCCTGACAGGTACGATTGCTGG-3' |
| | | 5'-CTGCAGGAGGGTTTCCTGTTGTCCAGAGTGTAGCGC-3' |
| | | 5'-CAAGGACATCTTTCCGGGACCGTGGTGTGAGTCCTC-3' |
| | | 5'-TTAAGGTCCGCTGGGATCTCTATGTGGCTGGCCATCAG-3' |
| | | 5'-TAAACGCTGACTTTGCTCTGCTTCCTCTGCGGGTTC-3' |
| | | 5'-CTCTCTGGGGAGTCACTCATGTCGCTGAGCTCCTTG-3' |
| | | 5'-CTGACAGTCCTCGGGACATGAAGCTGCACTTCCTGG-3' |
| *Pleurodeles waltl* | *Shh* | 5'-GGAGTAAGCTTCTTGGGGCGTCTCCTCTGGCCAATG-3' |
| | | 5'-CTTCTCGGCCACGTTGGGGATGAACTGCTTGTATGC-3' |
| | | 5'-GAGTTGCGCGTGATCTTGCCCTCATAACGTCCACTG-3' |
| | | 5'-GGGTTGTAATTAGGAGTTAGCTCCTTGAAGCGCTCC-3' |
| | | 5'-CGCTCCCGTGTTCTCCTCGTCCTTAAATATAATGTCA-3' |
| | | 5'-TCAGCTTATCCTTACACCTCTGGGTCATCAGCCTATC-3' |
| | | 5'-GACTCCAGGCCACTGGTTCATCACCGAGATTGCCAG-3' |
| | | 5'-TCTGAGGTGGTGATGTCCACTGCCCGACCCTCATAG-3' |
| | | 5'-CTCAAAGTAGACCCAGTCGAAGCCAGCCTCCACAGC-3' |
| | | 5'-CTCTGCTTTCACTGAGCAGTGGATGTGGGCCTTGGA-3' |

| | | |
|---|---|---|
| | | 5'-TGGGAAGCAACCTCCCGATTTTACAGCTACTGAGTT-3' |
| | | 5'-CAAGTCCTTCACGGGAATCCTCACCCCTTGCTCCAG-3' |
| | | 5'-AAGTCGCTGTAAATCAGCCTGCCCTCAACGTCCACG-3' |
| | | 5'-CTTTTCTGACCGTCTCTTCCTTGTCCATGAACAAGAGA-3' |
| | | 5'-TCTCCCGAGGCAGGGAGGTCTCTATCACGTAGAAGA-3' |
| | | 5'-TTTCCTGGGTGCTCTTGGGCTACAAAGAGGAGGTGG-3' |
| | | 5'-CTCCAGGTACACTCGATCCACCGTGGCTTCCCTTAG-3' |
| | | 5'-TAACAAGAGGCCAGCACCCTGTCTATGACCACGGTC-3' |
| | | 5'-ACAAAATACCGAAGCCCACTCTCAGAGGGGCGAAGG-3' |
| | | 5'-GGGAATGGCTGGAATAGTCTTGGGGAGAGAAGAATG-3' |
| | | 5'-CTCTGAGTACCAGTGGACTCCCTCTGCCTGAGAGGG-3' |
| | | 5'-CGCCTGTAACACCCATGTCCCTATCCGGTAGAGGAT-3' |
| *Xenopus laevis* | *prrx1* | 5'-CCTGTCCAGAAGGTGGTGGTTATAGCTGGAGGTCAT-3' |
| | | 5'-AATTGTCCAGACTGGCCGTGATGGGACTCTCCAAGC-3' |
| | | 5'-CCAGATCCAGCAGGTGACTCACCGAGAAGTTCTTTT-3' |
| | | 5'-CAGTAAGCTCCTGCCGGCTTCCCCTGATCCATCTTC-3' |
| | | 5'-TTGCTGAGGAGTGTCACTTCCACTGGTCAAACCCGG-3' |
| | | 5'-TCCTTTGCTTTCTCTTCTTCTTCTGTTCTGCATTCATCTGT-3' |
| | | 5'-GAGCTTGCAGTTGACTGCTGTTAAATGTAGTTCTGTTCC-3' |
| | | 5'-GCATCTGGATAATGCGTCCTCTCAAACACTCTTTCCA-3' |
| | | 5'-AGGTTAACTCTGCGCGCTAGGTCTTCCCGTACAAAG-3' |
| | | 5'-CTGTTCTGGAACCAAACCTGGACTCTGGCTTCGGTG-3' |
| | | 5'-AGCATGGCTCTTTCATTTCTGCGGAACTTGGCTCTT-3' |
| | | 5'-GCATAGGATTTGAGAAGGGAAGCATTCTTGTTCGCC-3' |

| | | |
|---|---|---|
| | | 5'-GGTACAATGGGCTGTTCCACAGCGGTCACATCTCCT-3' |
| | | 5'-CTGTGCCCCAAGATAGATACTCATTGGGCCTCGGGG-3' |
| | | 5'-TAGAGGAATAGCTAGCCATGGCACTATATGGGGAGG-3' |
| | | 5'-TAAGGTTGGCGATGCTGTTGGCCATGTTCATCCCCT-3' |
| | | 5'-TGGTTCCTCTGTAAACTGTATTCCTTTGCCTTCAGCC-3' |
| *Xenopus laevis* | *sox9* | 5'-TGCTCTTCTGACATCTTCATGAAGGGATCCAAGAGATTCAT-3' |
| | | 5'-TGTTTTCTTGGGGTCTGGTGTTCTCCGTGTCGGAGC-3' |
| | | 5'-TCTCCTTCTTCATCTCCTGGTCCCCTTTGGGGAAAG-3' |
| | | 5'-CTTCTCTGATGCACACGGGGAACTTCTCGTCCTCTG-3' |
| | | 5'-CAGGGTCCAGTCATATCCCTTCAGCACCTGGCTGAC-3' |
| | | 5'-CTTGCTGGATCCATTAACTCTGACTGGCATCGGTAC-3' |
| | | 5'-GAAGGCATTCATGGGTCTCTTGACATGGGGCTTGTT-3' |
| | | 5'-GATATTGGTCTGCCAGCTTCCTCCTTGCAGCCTGTG-3' |
| | | 5'-CCAGCGTCTTGCTGAGTTCTGCATTGTGCAGATGGG-3' |
| | | 5'-GCGTTTCTCACCTTCATTCAAAAGCCTCCATAACTTCC-3' |
| | | 5'-GTTGGACCCTCAGCCTCTCTGCTTCCTCCACGAAAG-3' |
| | | 5'-GTGGCTGATACTTGTAGTCGGGATGATCCTTCTTGT-3' |
| | | 5'-GTTCTGACTGCCCGTTCTTAACAGACTTTCTGCGCC-3' |
| | | 5'-TAGGGGAGATGTGGGTCTGCTCAGCACTATCATCTT-3' |
| | | 5'-GTGGGGAGTCAGCCTGTAGGGCCTTGAAAATTGCAT-3' |
| | | 5'-CCAGGAGAGTGGACTTCACTCATGCTGGAAGTAGAAT-3' |
| | | 5'-GGAGTCGGTGGGCCTTGGGATTGACCTGAATGTTCT-3' |
| | | 5'-AAGTCTGGCTTTCCAGGCTGGACATCTGTCTTGGGG-3' |
| | | 5'-GAAATCAATGTGAGGCGGTTGCCTACCGCTCTCCTG-3' |

| | | |
|---|---|---|
| | | 5'-GACCTCACTGCTCAGCTCACCAATGTCCACATCACG-3' |
| | | 5'-GGTCAAATTCATTGACGTCAAAGGTTTCGATGGTGGAGAT-3' |
| | | 5'-TAGGGGTGCTGCTGATGCCATAACTGCCTGTGTACG-3' |
| | | 5'-TAGACATCCAGGCAGAACCAGCACCTGTAGTGGCAC-3' |
| | | 5'-TTGACAGTGAGTGTTGCTGAGGCTGCTGTTGCTGTT-3' |
| | | 5'-TCCTTTGCTGGGACTGGCTTTGCTCGCTGTTTATGG-3' |
| | | 5'-AATGACTTGGGCTCAGTTGCTCAGTCTTGATGTGTG-3' |
| | | 5'-CTGTAGGTTGAAGGAGCTGTAGTTCAGTTGCTGGGG-3' |
| | | 5'-TGCCCGGGTGATGGTTGGGTAGGAAGAGCTGTAATG-3' |
| | | 5'-GGAGTTGGAGCCCTGGTGCTCTGTGTAGTCATACTG-3' |
| | | 5'-GAGTAGAGACCAGAATTTTGACCACTTGCGTGAGTGTAAT-3' |
| | | 5'-ATGGGGCGTTGGCTTGGATTCATGTAGGTAAAGTTG-3' |
| | | 5'-GATGGAACTCCCGTTGTGTCTGCAATAGGGGTGTAC-3' |
| | | 5'-AGTTGTGTATAGACAGGCTGCTCCCAGTGCTGTGGG-3' |
| *Mus musculus* | *Fgf8* | 5'-CCTCGTACTTGGCGTTCTGCAGCGCCGTGTA-3' |
| | | 5'-AAACCCAACAGCAAACAATATGCACAACTAGAAGGCA-3' |
| | | 5'-GCTGGTGCGGCTGTAGAGCTGGTAGGTCCG-3' |
| | | 5'-CTGCGCGTGAAGGGCGGGTAGTTGAGGAAC-3' |
| | | 5'-GCGGCTGAGCTGATCCGTCACCAGGCTCTG-3' |
| | | 5'-TAGTTTGGGGCTGGATTTGTAAAGTTCTTTCCAGCCC-3' |
| | | 5'-CGGCCCTTGCGGGTAAAGGCCATGTACCAG-3' |
| | | 5'-CCTTCTTGTTCATGCAGATGTAGAGACCTGTCTCTGC-3' |
| | | 5'-GACAAGCCGAAGGTGCGGAGGCTGAGGGAG-3' |
| | | 5'-CAGGCGCTTCATGAAGTGCACCTCGCGCTG-3' |

| | | |
|---|---|---|
| | | 5'-GTCTCCGTCTTCTGCCATGGCGTTGATGCG-3' |
| | | 5'-CTTCTGCGCAGCGCTCCTTACCTGGGCTTG-3' |
| | | 5'-GGTCCTCGTGTCCCTGCCCGAGAGTGTCAG-3' |
| | | 5'-GCCGCGAACTCGGACTCTGCTTCCAAAAGT-3' |
| | | 5'-CGGCTATCCCGACGCGGCTCGTGGAATGTC-3' |
| | | 5'-TGTCTCTCTGTTCCTTAGATCCCTCCTCGGGG-3' |
| | | 5'-CTTGTTGGCCAGGACCTGCACGTGCTTCCC-3' |
| | | 5'-GTTGTTCTCCAGCACGATCTCTGTGAATACGCAGT-3' |
| *Mus musculus* | *Fgf10* | 5'-TGTGAAGTCTTCCGAAAGCCACTTCTTGTTTGGGG-3' |
| | | 5'-GTCTCTTTGGAGTTGTCAGAACTGGAGGCAGCTGC-3' |
| | | 5'-TCTCTGCAGAGTCCCAGCCTGCTGGCCACTTAAAA-3' |
| | | 5'-CAGGACGGTGAACAAAACCCAAAGTGGGTGGGGTG-3' |
| | | 5'-ACTCCTTTCTTCACCATGTTAGATGCAAAAGAGGTCTGAAA-3' |
| | | 5'-GAACTGGTGGTGTCGGTCACCAGAGCTCTTCGCTC-3' |
| | | 5'-TAACTTCTAGGAAGGACCGGCTGCCTGTCCTCGCTC-3' |
| | | 5'-TACGGATCTGGCCAGAAGCGAGTGCACCAACATCC-3' |
| | | 5'-ACATACTGGAAGGGTAAGACCTGCTGCGAGGCAGA-3' |
| | | 5'-AACTCTCGGCACTGGAAATTGTCTCATCAGAAGGA-3' |
| | | 5'-AGGCACAATGTGTCAGTATCCATTTCCACATTGTACTGA-3' |
| | | 5'-AGCTTGGCAGGTGACAGGGAACGAAGACACCAAAA-3' |
| | | 5'-CTTGGAGGTGATTGTAGCTCCGCACATGCCTTCCC-3' |
| | | 5'-GGTGAAGGAGAACAGCCTTCTCCAGCGGACATCTC-3' |
| | | 5'-TGACCTTGCCGTTCTTCTCAATCGTGAGAAAGTACTT-3' |
| | | 5'-CTGTTGATGGCTTTGACGGCAACAACTCCGATTTC-3' |

| | | |
|---|---|---|
| | | 5'-TGAGCCATAGAGTTTCCCCTTCTTGTTCATGGCTAAGTAAT-3' |
| | | 5'-GCTGCCAGTTAAAAGATGCATAGGTGTTGTATCCATTTTCC-3' |
| | | 5'-TCCATTCAATGCCACATACATTTGCCTGCCATTGT-3' |
| | | 5'-GATCGTCATGGGGAGGAAGTGAGCAGAGGTGTTTT-3' |
| | | 5'-GGTTGTACTGCATCCACCAACAGTGTTTTCTTCTATGTTTG-3' |
| *Mus musculus* | *Gdf5* | 5'-GTCTGTATCCAGTCCCATAGTGGAAGTGCTCCCTTTCATCATCAT-3' |
| | | 5'-TGTGGATTCAGGGTCCATAGAGTTCATTAGGGTCTGATTTCATCATCAT-3' |
| | | 5'-GAAAATAACTCGTTCTTGAAAGGAGAAAGCCGACCGCTTTCATCATCAT-3' |
| | | 5'-TCTTTAGTCCAGATCACATCCAGAGAGTAAGCCTGCTTTtCATCATCAT-3' |
| | | 5'-CTGTACAGGGAGAGCATGTATTCGTGGGGTGTTTtCATCATCAT-3' |
| | | 5'-AAACACTAGGAACAAGGCTTTCTCGTGGACCTGTTTtCATCATCAT-3' |
| | | 5'-AAGGTGATTCAAGGAGGAGACAGCTCTCCCTTAAAGATTTCATCATCAT-3' |
| | | 5'-CCACGACCATGTCCTCGTACTGTTTATACACCACGTTTCATCATCAT-3' |
| | | 5'-GGGGCATCTTTCTGGGTTCATCCTTCTTGGCTTTCATCATCAT-3' |
| | | 5'-TCTAATTAGGCTCATACTCTTCTCTTCACCCCTGCCCTTTCATCATCAT-3' |
| | | 5'-CCAGCCGCTGAATGACACCACAGAGAAAGGTTTCATCATCAT-3' |
| | | 5'-CAGTGGAAGGCCTCATACTCAAGAGGTGCGATTTCATCATCAT-3' |
| | | 5'-AATTTCGGAAGAGCTTCCAGATGTCGAACACCTCCTTTCATCATCAT-3' |
| | | 5'-AACTTCTCTGGAGAGTCTGGAGAGAAATGAAGAGGCCTTTCATCATCAT-3' |
| | | 5'-TTCATACACAGTCTTGTCATCCTGGCCAGAGCGTTTCATCATCAT-3' |
| | | 5'-CCGGGCAGGAGAGATGGTGAACTGAGTCTCTTTCATCATCAT-3' |
| | | 5'-ATTAAGAACACCTTTCCTGAGCCCCAGGCCCTTTCATCATCAT-3' |
| | | 5'-GATGTCAAACACGTACCTCTGCTTCCTGACCGCTTTCATCATCAT-3' |
| | | 5'-CCTCCCTTTCTGTCAGCATCGGACAGCGTCTTTCATCATCAT-3' |

5'-GAGCGCAAGGGGAACTCACACAGTCCTTCGTTTCATCATCAT-3'
